# Rare sex but a long life sustain seaweed populations at the warm edge of their range

**DOI:** 10.1101/2025.06.23.660995

**Authors:** Sofie Vranken, Frederique Steen, Sofie D’hondt, Marc Verlaque, Conxi Rodriguez-Prieto, Christophe Viera, Thierry Thibaut, Kenny Bogaert, Olivier De Clerck

## Abstract

**Aim:** The life cycle of many organisms is all but stable across their distribution range. Most commonly, populations respond to environmental variation by shifting the timing of reproductive events (phenology). Or more profoundly, populations may (partly) shift their mode of reproduction from sexual to asexual. Life cycle variation can impact reproductive success, gene flow, genetic diversity, and, ultimately, the evolutionary trajectory of populations. Understanding the factors that influence life cycle variation is essential for grasping the biology and ecological roles of species. This study investigates the variation in life cycles and its effects on the genetic diversity of a brown seaweed, *Dictyota*, across its European range.

**Location:** North-East Atlantic Ocean and the Mediterranean Sea.

**Taxon:** *Dictyota dichotoma* (Hudson) J.V. Lamouroux (Phaeophyceae, Dictyotales)

**Methods:** We monitored phenology, fertility and lifespan in Atlantic and Mediterranean populations at its northern and southern boundaries, and used microsatellite markers to assess how these factors influence genetic and genotypic diversity.

**Results:** We observed significant differences in phenology, reproductive strategies, and genetic diversity among northern and southern European populations. In Mediterranean populations, *D. dichotoma* exhibited sporophytic dominance with gametophytes being extremely rare, suggesting a shift towards asexual reproduction. In contrast, North-East Atlantic populations displayed more pronounced seasonal reproductive patterns with higher frequencies of gametophytes, indicating predominant sexual reproduction. However, genetic analysis showed lower allelic richness and unique alleles in northern populations, whereas southern populations were genetically more diverse, reflecting historical biogeographic processes. Clonal reproduction was more pronounced in the Mediterranean populations, influencing the spatial genetic substructure and contributing to lower genotypic diversity compared to Atlantic populations.

**Main conclusions:** Our findings demonstrate that life cycle variation and phenology in *D. dichotoma* are closely tied to regional environmental conditions and have significant implications for population structure and genetic diversity. Results highlight how shifts in reproductive strategies contribute to the evolutionary and ecological dynamics of marine macroalgae across biogeographical gradients.

## Introduction

An organism’s life cycle is a crucial factor that underpins fundamental ecological and evolutionary processes. By influencing reproduction, survival and genetic diversity, the life cycle directly influences the fitness of a population and, therefore, whether populations will thrive or perish under certain conditions. Species, but also populations of the same species, can exhibit significantly different life cycle variations. Understanding the characteristics that influence life cycle variation is fundamental to comprehending species’ biology and functionality within an ecosystem. Importantly, documenting life cycle variation and its effects on population dynamics across a species’ range may inform us on the faith and persistence of populations under future climatological scenarios.

A life cycle can include different modes of reproduction, i.e. sexual or asexual reproduction or both. Although sexual reproduction (meiotic sex sensu Stacy A. Krueger-Hadfield (2024)) comes with a higher energetic and genetic cost (only half of a parent’s genes are transferred to the next generation), it is generally considered more beneficial as it includes genetic recombination, which increases genetic and genotypic diversity in the progeny, favouring adaptation. In asexual reproduction (vegetative propagation or apomixis sensu Stacy A. Krueger-Hadfield (2024)), where there is no genetic recombination, genotypic diversity is expected to decrease. Furthermore, selection efficacy is expected to decrease through higher linkage among alleles and selection interference (Otto, 2021), ultimately leading to accumulations of deleterious alleles and decreased fitness (Muller, 1964). Yet, multiple forms of asexual reproduction are present across a myriad of species, often at the edge of species geographical distribution where environmental conditions reduce the relative benefits of sexual reproduction and promote asexual reproduction (Silvertown, 2008). Whether a species reproduces asexually or sexually has profound effects on its genetic diversity and evolutionary trajectory (Hartfield, 2016; Vranken, Scheben, Batley, Wernberg, & Coleman, 2022), though the relative consequences of asexual versus sexual reproduction are not yet understood for many natural populations.

Phenology, the timing of reproductive events, influences population fitness and genetic diversity by influencing, amongst others, reproduction success and gene flow with surrounding populations (Forrest & Miller-Rushing, 2010; Katal, Rzanny, Mäder, & Wäldchen, 2022). Phenology is known to be greatly affected by environmental conditions and it has emerged as one of the most common biological reactions to climate change across a wide range of species (Chmura et al., 2019; Cleland, Chuine, Menzel, Mooney, & Schwartz, 2007; Parmesan, 2006). Changes in phenology can be either beneficial or detrimental, depending on the specific ecological and evolutionary context. As changes in phenology can differ in extent and direction among organisms, asynchrony among species or populations can reduce gene flow and a population’s genetic diversity (Edwards et al., 2024). Even though attention to documenting and forecasting the impacts of climate change has increased interest in phenological research (Forrest & Miller-Rushing, 2010; Katal et al., 2022), it still remains an overlooked aspect of many, less studied, groups of species (de Bettignies, Wernberg, & Gurgel, 2018; Stacy A. Krueger-Hadfield et al., 2023).

Seaweeds, and brown algae in particular, are a group of habitat-forming organisms that form a substantial part of our sub- and intertidal ecosystems (Bringloe et al., 2020), that are renowned for their large variety of life cycles (Bell, 1997; Heesch et al., 2021). Their sexual life cycle is predominantly characterised by a haplodiplontic life cycle where there is an alternation of a diploid sporophyte that produces haploid spores after meiosis, which subsequently grow out to a haploid gametophyte. The gametophyte produces male and female gametes through mitosis, which after fertilization, the resulting zygote develops into a diploid sporophyte (Vranken et al., 2023). Spore and gamete release and maturation are regulated by a combination of environmental cues, including photoperiod, temperature, nutrients and water movement but also the presence of female pheromones (de Bettignies et al., 2018) or lunar cycles (Müller, 1962). For example, the brown kelp *Undaria pinnatifida* (Laminariales) shows pulses in recruitment in California that coincide with temperature drops and nutrient pulses (de Bettignies et al., 2018), indicating that temperature and/or nutrient levels might regulate the sexual life cycle by influencing spore maturation and/or release when seawater temperature or nutrient levels change vastly or abruptly.

Yet, many brown seaweeds are also capable to reproduce asexually through a variety of mechanisms (Coelho et al., 2007; Vranken et al., 2023). For example, in some species (e.g. *Ectocarpus sp., Laminaria sp.*) the haploid cells of the gametophyte can restore diploidy without fertilisation but by endomitosis (Bothwell, Marie, Peters, Cock, & Coelho, 2010; Goecke, Klemetsdal, & Ergon, 2020; Müller, 1967; Oppliger et al., 2014; Shan, Pang, Wang, & Li, 2021). Next to this, apomeiosis, where meiosis in the sporophyte is replaced by a mitosis, resulting in diploid spores that develop into a new sporophyte are also observed (Bothwell et al., 2010; Goecke et al., 2020). Lastly, many brown algae can reproduce by fragmenting either haploid or diploid multicellular forms, bypassing the sexual cycle entirely (Ardehed et al., 2015; Bogaert, Delva, & De Clerck, 2020; Vranken et al., 2022). Many seaweeds produce asexually at the edge of their species distribution driven by environmentally challenging conditions. For example, *Scytosiphon lomentaria* (Ectocarpales) switches to an asexual life cycle in colder waters (Hoshino et al., 2021) and *Fucus radicans* (Fucales) tends to produce asexually when salinity is low (Ardehed et al., 2015). Such marginal populations are often characterized by increased genetic isolation, genetic differentiation, and dominance of one ploidy level (Stacy A. Krueger-Hadfield, 2020), which can impact individual and population performance. Understanding how environmental gradients influence reproductive modes and genetic diversity in seaweeds is crucial, especially under scenarios of rapid climate change.

Here, we investigate life cycle variation and its consequences on the genetic diversity of *Dictyota dichotoma* (Hudson) J. V. Lamouroux (*Dictyota* hereafter, Dictyotales). In Europe, the species ranges from the Canary Islands to the southern Norwegian coast, including the Mediterranean (Tronholm et al., 2010). In the North East Atlantic, macroscopic thalli disappear in winter, while in the Canary Islands they are present almost all year round (Tronholm, Sanson, Afonso-Carrillo, & De Clerck, 2008). How Mediterranean populations that experience stressful summer temperatures respond and how this affects genetic diversity is the main focus of this research. Clonal propagation through fragmentation has been observed in other *Dictyota* species (Herren, Walters, & Beach, 2006), as well as the ability to regenerate the sporophyte phase through the production of sporophytic *in situ* germlings (Bogaert et al., 2020; Hwang, Kim, & Lee, 2005). As temperature plays a prominent role in the control of sporogenesis and growth in this species (Bogaert, Beeckman, & De Clerck, 2016), we assess here whether variations in the reproductive phenology and reproduction mode can be linked to sea surface temperature and how this affects the genetic diversity. For this, we develop co-dominant single-locus microsatellites to screen for patterns of asexual reproduction and estimate the population genetic diversity in 14 populations spanning almost its entire species distribution in Europe. We then monitor the life stage (gametophyte/sporophyte) and the reproductive state (phenology) of four populations to determine whether the Mediterranean populations are dominated by sporophytes year-round and how this differs from Northern Atlantic populations. Next, we investigate the spatial scale of clonality, small-scale population structure and the life span of three Mediterranean populations to test whether the lack of sex can be compensated by perenniality. In addition, we assessed whether *Dictyota* could reproduce asexually by apomeiosis by assessing spore ploidy using the microsatellite markers.

## Materials and Methods

### Phenology

To examine phenological differences among populations of *Dictyota*, two Mediterranean populations (Calanque de Sormiou at Marseille and Carry-le-Rouet, Bouches-du-Rhône, France) and two NE Atlantic populations (Wimereux, Pas-de-Calais, France and Goes, Zeeland, the Netherlands) were monitored and sampled on a monthly basis from May 2012 to May 2013 (Fig. 1) (Steen, Verlaque, D’hondt, Vieira, & De Clerck, 2019). At each location, we sampled 50 individuals along a 50m transect, randomly selecting five individuals every 5m. The ploidy level of each individual was scored as haploid when male or female gametangia were visible and as diploid when sporangia were present. When not reproductive, individuals were scored as sterile. To investigate variations in fertility in relation to local thermal regimes, mean monthly sea surface temperatures for each location were obtained from the OCLE Sea Surface Temperature (SST) dataset (de la Hoz et al., 2018). To quantify regional differences in thermal performance of fertility independently of seasonality, we modelled temperature-dependence of the proportion of reproductive individuals according to (Rodríguez, García, Carreño, & Martínez, 2019): Fertility (T) = *a*/[[1 + exp(-*b*\*(*T*-*c*))] * [1+exp (*d*\*(*T*-*e*))]] where *T* is the temperature (°C), *a* the maximum fertility, *b* the slope of the ascending part of the curve, *c* the temperature value for the midpoint of the ascending sigmoid, *d* the slope of the descending part of the curve, and *e* the temperature value for the mid-point of the descending sigmoid. To capture regional variation, we tested all combinations (2⁵) of these parameters being either shared across regions or region-specific. Models were fit using nonlinear least squares (nlsLM), with starting values: *a* = 0.5, *b*, *d* = 0.1, and *c*, *e* = mean temperature; bounds were set as: *a* ∈ [0.01, 1], *b*, *d* ∈ [0.01, 2], *c*, *e* ∈ [0, 30]. Model performance was evaluated using the Akaike information criterion (AIC), Bayesian Information Criterion (BIC), and a pseudo-R² (1 – RSS/TSS), and the best Rodriguez model was selected based on the lowest BIC (-26.15) and AIC (-40.77). To compare its performance, we also fit alternative models: a linear non-temperature dependent model, Brière model (Briere, Pracros, Le Roux, & Pierre, 1999); Blanchard model (Blanchard, Guarini, Richard, Gros, & Mornet, 1996) with comparable region-specific structure and evaluated using the same criteria to identify the best-fitting representation of thermal performance.

**Figure 1.**
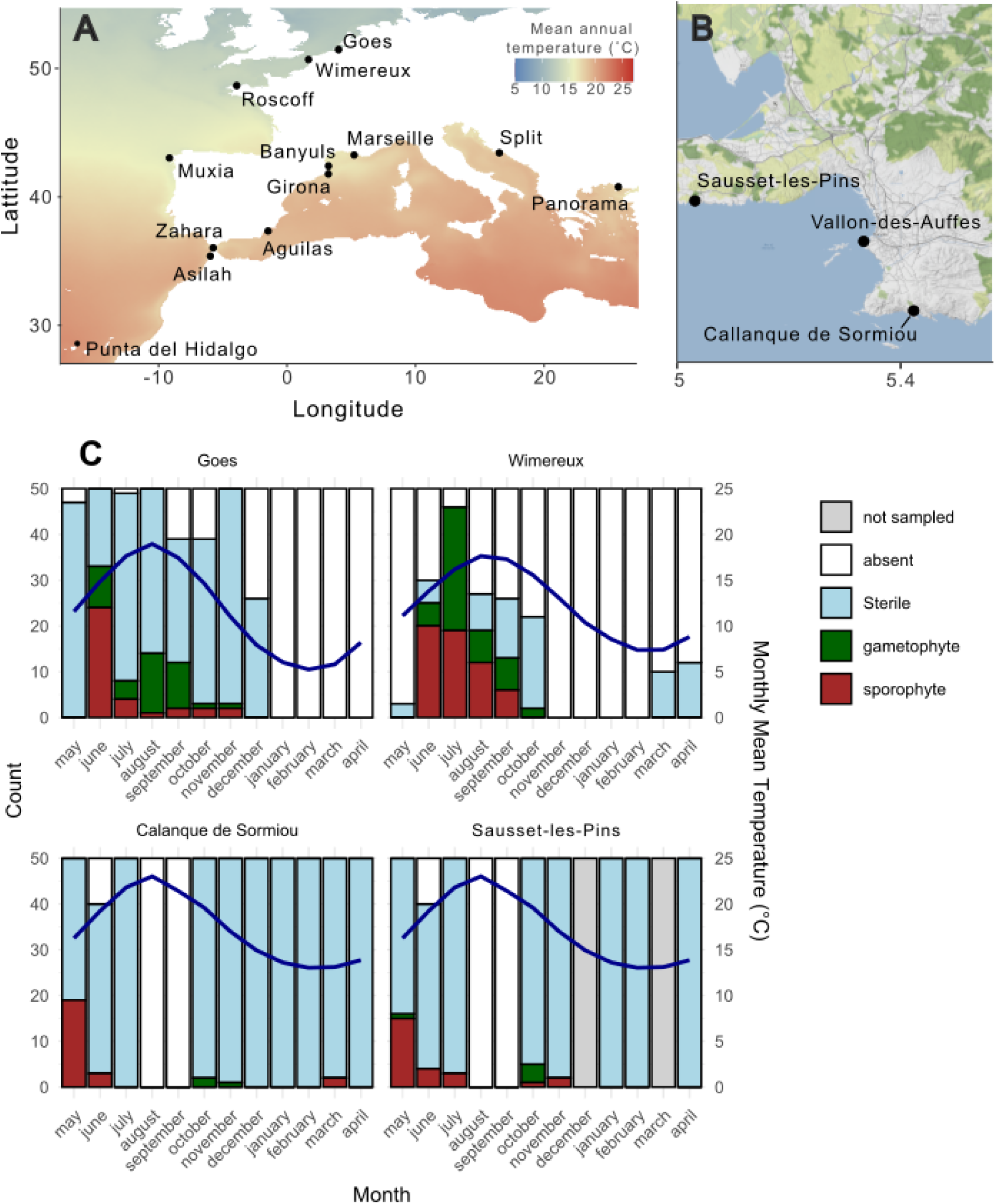
A) Sampling locations of *Dictyota dichotoma* in Europe, B) Detailed map of the sampling locations in the Marseille area, C) Proportion of sterile (blue), fertile gametophyte (green), fertile sporophyte (purple) and non - sampled (grey) individuals (on a total of 50 individuals) throughout the year in two Atlantic populations (Goes and Wimereux) and in two Mediterranean populations (Sausset-les-Pins and Callanque de Sormiou). The blue line represents the local mean monthly sea surface temperatures.

### DNA extraction and genetic & genotypic diversity assessment

To assess genetic diversity and test for the presence of clonal reproduction in 13 *Dictyota* populations across its native range, we used genetic data from 2015-2016 (Steen, Verlaque, D’hondt, Vieira, & De Clerck, 2019) for microsatellite analysis. These populations included European Atlantic coasts (European Atlantic, Strait of Gibraltar, and Macaronesia) and the Mediterranean (Northwestern Mediterranean, Adriatic Sea, and Eastern Mediterranean). Where possible, we aimed at collecting a minimum of 20 sporophytes separated at least 1m from one another and a few gametophytes where possible (to test the presence of null alleles, see further). Apical tips were sampled and stored in silica for DNA analysis. Genomic DNA was extracted using an in-house CTAB method following Steen, Vieira, Leliaert, Payri, and De Clerck (2015). All samples were genotyped using microsatellites as described by Vranken, Steen, D’hondt, and De Clerck (submitted). Allele sizes were manually scored using the GENEIOUS Prime microsatellite plugin (2023.2.1) (https://www.geneious.com), and the ploidy of the samples was assessed as above. The frequency of null alleles was directly estimated from the non-amplified samples of the haploid gametophytes, disregarding technical errors. Given the very low proportion of non-amplification (maximum 1% for locus Dd 1 and Dd7) of known haploid samples, we considered that there was no effect of null alleles. The absence of large allele dropout and stutter alleles was confirmed for the sporophyte data using MICROCHECKER (Van Oosterhout, Hutchinson, Wills, & Shipley, 2004). One locus (DdB) was removed due to putative allele stuttering. No more than 10% missingness per sample and 5% per locus were allowed, giving a total missingness of 0.22%. As the number of gametophyte samples was not large enough, diversity indices were only calculated on the sporophyte subpopulation.

Allele sizes were manually scored using the GENEIOUS Prime microsatellite plugin (2023.2.1) (https://www.geneious.com), and the ploidy of the samples was assessed as above. The frequency of null alleles was directly estimated from the non-amplified samples of the haploid gametophytes, disregarding technical errors. Given the very low proportion of non-amplification (maximum 1% for locus Dd 1 and Dd7) of known haploid samples, we considered that there was no effect of null alleles. The absence of large allele dropout and stutter alleles was confirmed for the sporophyte data using MICROCHECKER (Van Oosterhout, Hutchinson, Wills, & Shipley, 2004). One locus (DdB) was removed due to putative allele stuttering. No more than 10% missingness per sample and 5% per locus were allowed, giving a total missingness of 0.22%. As the number of gametophyte samples was not large enough, diversity indices were only calculated on the sporophyte subpopulation.

To assess clonality, the number of multilocus genotypes (MLGs) was calculated using the R-package ‘poppr’ (Kamvar, Tabima, & Grünwald, 2014) for the total sample size and rarified on a sample size of 20. Only samples with repeated MLGs, where the probability that they do not come from separate sexual reproduction events (psex < 0.05), were considered as samples from the same MLG or as clones. On the contrary, when psex > 0.05, samples with repeated MLGs were not considered clones but as belonging to different MLGs. Clonal richness (R) was calculated as R = (G-1)/(N-1) with G as the unique number of identified MLGs and N as the total number of samples (Arnaud-Haond, Duarte, Alberto, & Serrao, 2007; Dorken & Eckert, 2001). The distribution of the clonal memberships was described with the Pareto Beta index (high B value for high evenness, low B value for skewed distribution with the dominance of a few MLGs). Genetic diversity within sampling sites was assessed by calculating observed heterozygosity (Ho), expected heterozygosity (He) and the inbreeding coefficient (F_IS_) with the R package ‘Hierfstat’ (Goudet & Jombart 2020). Allelic richness standardised for sample size was calculated with the R package ‘PopGenReport’ (Adamack & Gruber, 2014). Significance of F_IS_ from zero was calculated with 9999 permutations. The variance and the distribution patterns of F_IS_ values across loci were also evaluated since these are more informative compared to average F_IS_ values in cases of partial clonality (Reynes et al., 2021; Stoeckel, Porro, & Arnaud-Haond, 2021). Linkage disequilibrium (LD) between among locus pairs was assessed for each population with the ‘Genepop’ R-package (Rousset, 2008). p-values were adjusted for multiple comparisons with the Bonferroni correction. We reported the pairwise linkage disequilibrium as the number of significant pairwise LD per total number of comparisons per population. Significant multi-locus linkage disequilibrium was estimated using the Standardized Index of Association (rbarD) implemented in the R-package ‘poppr’ (Kamvar et al., 2014) and performing 999 permutations. These statistics were calculated for both the entire dataset and the clone-censored dataset.

To test if there was a link between genetic or genotypic diversity and temperature, the correlation and its p-value were calculated between average yearly temperature and expected heterozygosity, allelic richness, clonal richness, Pareto B, and the variance of F_IS_. Pearson correlation was used for normal-distributed data (allelic richness), and Spearman rank correlation was applied for non-normal-distributed data.

### Fine-scale clonal and spatial structure

Fine-scale clonality and spatial structure were assessed in Marseille in 2021 at three locations (Sausset-les-Pins, Callanque de Sormiou & Vallon des Auffes) with a 50cm x 50cm grid. Two quadrats were placed 10 cm apart, and per quadrat, six individuals of *Dictyota* were collected if present. This was repeated over a transect parallel with the coast, with one, 10, and again one meter between quadrats. Per location, this was repeated at two sites +-100m apart. These samples were processed for microsatellite analyses as above. No more than 10% missingness per sample and 5% per locus were allowed, resulting in a dataset of 223 samples (48 quadrats) genotyped for twelve loci with a total missingness of 0.6%. The absence of null alleles, large allele dropout and stutter alleles was confirmed with MICROCHECKER. MLGs were defined as above. First, we visualised within-population clonal structure by plotting the repeated MLGs as a heatmap for each grid cell and mapping the genets on the grid. This allows us to estimate and visualise the minimum area a repeated MLG can occupy. Next, a spatial autocorrelation was performed to assess the scale of spatial dependence of the genetic and genotypic diversity using SPAGEDI 1.5 (Hardy & Vekemans, 2002). We estimated Ritland’s kinship coefficients (Ritland, 1996) among pairs of individuals belonging to six distance classes ranging from 0.5 to 250m. Distance classes were selected to have > 100 pairwise comparisons, a participation index > 50% and a coefficient of variation ≤1 (Hardy & Vekemans, 2002). Significance from zero was tested for every distance class by comparing the observed Ritland’s coefficients to the frequency distributions calculated by 999 permutations. Spatial autocorrelation was performed per location on all individuals (ramet level, including repeated MLGs) and only with unique genetic strains (genet level, repeated MLGs removed).

### Perennial persistence vs annual recruitment

To assess whether Mediterranean populations of *Dictyota* could survive by perenniality, either by survival of the upright thallus or by sprawling from a cryptic perennial base, we monitored four sites (Sausset-les-Pins, Calanque-de-Sormiou, La Fosca & S’Alguer) at two locations (Marseille, France, and Girona, Spain) for two consecutive years (2022-2023). For this purpose, a 50 x 50 cm grid quadrat was placed on the reef, and the exact location of the quadrat was marked by drilling or glueing markers into the reef. Per quadrat, we sampled the upper part of all *Dictyota* individuals present, leaving the base of the individuals intact so that regrowth was possible. Per sampled individual, grid coordinates were noted. Depending on the availability of *Dictyota*, one to four quadrats were marked (5-100m apart) per site. All samples were processed for microsatellite analysis. The absence of null alleles, large allele dropout and stutter alleles was confirmed with MICROCHECKER (Van Oosterhout et al., 2004) and all 12 loci were retained. After filtering out samples and loci with more than 10% and 5% missing data, we retained a dataset with a total missingness of 1.1% and 377 individuals. Ploidy levels were again based on the genotypes and used to distinguish sporophytes from gametophytes (see above). We identified the number of repeated MLGs and assessed whether these remained present over multiple years and occurred in the exact locations.

This analysis was repeated for the sporophytes, including the samples from Marseille used for the fine-scale hierarchical structure, which were sampled in 2021 in the same Marseille region but not in the same grid quadrats. This allowed us to identify if MLGs can be retained over multiple years on a broader geographic scale. The combined dataset was filtered as above, and included 580 sporophytes with a total missingness of 0.75%.

### Tetrad analysis

To exclude the presence of asexual reproduction through apomeiosis as reported for other *Dictyota* species (Hwang et al., 2005) and confirm the production of four haploid meiospores in the sporophyte, we isolated cultured tetrads (spore quartets) from four independent *D. dichotoma* sporophytes: UGCC0031 (Goes, Atlantic Ocean), UGCC0023 and UGCC0019 (Marseille, Mediterranean Sea), and UGCC0003 (Croatia, Mediterranean Sea). For each strain, the sex ratio of the offspring was scored based on their reproductive structures. Deviance from 50:50 male:female ratio was assessed for the offspring per parental sporophyte with an exact binomial test. All offspring were processed for microsatellite analysis as explained above. Ploidy levels of the offspring were assigned based on the genotypes in which samples with only homozygous loci were regarded as gametophytes and any samples with at least one heterozygous locus as a sporophyte (Heiser et al., 2023; Stacy A. Krueger-Hadfield et al., 2016).

## Results

### Phenology

A total of 1,425 specimens were sampled at two southern (Calanque de Sormiou, Sausset-les-Pins) and two northern populations (Goes, Wimereux) to assess the phenology of *Dictyota*. Due to poor weather conditions, Sausset-les-Pins was not sampled in December and March. Macroscopic thalli were absent throughout August and September at the southern sites, when temperatures were highest (Fig. 1C). For the Atlantic populations, no *Dictyota* individuals were observed in Goes from January to April and in Wimereux from November to February when temperatures were lowest. The proportion of fertile individuals was generally lower in the southern populations compared to the northern ones. Out of 900 samples taken from the two southern populations, fertile gametophytes were extremely rare (N = 8). They were mostly observed in autumn: one male and three female gametophytes in October in Sausset-les-pins and two male and one female gametophyte in October and November for the Calanque de Sormiou population. In spring, only one fertile male was observed in May in Sausset-les-Pins. In contrast, the northern populations in Wimereux and Goes showed fertile gametophytes from June through October and November, respectively. Sporophytes were present from June to August for Wimereux and from June to November in Goes. In Wimereux, almost all individuals were fertile in June and July, and most of the individuals in Goes were fertile in June. In the southern populations, however, less than 40% of the individuals were fertile during the most fertile month (May).

Model comparison identified a thermal performance function with region-specific parameters (Fig. S1; Table S1) as the best-performing model (R^2^=0.54). The predicted thermal optima for fertility were 15.9 °C in the Mediterranean and 16.4 °C in the North Sea. Alternative formulations, including linear, Brière, Blanchard, and Rodriguez temperature response models, performed less well in terms of fit and explanatory power. These results indicate that temperature is a key explanatory variable for spatial variation in fertility. Despite pronounced differences in observed phenological patterns between regions, the model infers similar thermal optima, suggesting that divergent reproductive phenologies are driven by seasonal differences in temperature exposure rather than fundamental differences in physiology between Mediterranean and North Sea populations.

### Tetrad analysis

For each strain tested, the sex ratio of the offspring was scored based on their reproductive structures: UGCC0031 showed 55% male / 45% female (n = 40), UGCC0023 showed 43% male / 57% female (n = 7), UGCC0019 showed 56% male / 44% female (n = 16), and UGCC0003 showed 50% male / 50% female (n = 24) (Table S2). Male:female ratio of the offspring did not deviate from 50:50 for all three sporophytes tested. (Exact binomial test p > 0.05). All 87 genotyped offspring—32 from UGCC0031 (Goes - Atl. Oc.), 24 from UGCC0019 and UGCC0023 (Marseille), and 23 from UGCC0003 (Croatia – Med Sea)—were haploid, with no evidence of diploid individuals regardless of their origin. These results indicate the absence of apomeiosis and confirm that sporophytes in *Dictyota* produce four distinct haploid meiospores through regular meiosis.

### Genetic & genotypic diversity

We assessed genetic diversity and tested for the presence of clonal reproduction in *Dictyota* populations across its native range by using microsatellites. The number of rarified MLGs on a total of 20 individuals was lowest for Panorama (eMLG = 3) and Split (eMLG = 13), in the Eastern Mediterranean and the Adriatic Sea, respectively (Table 1). But also Asilah, where rarefying to 20 was not possible, had a low number of MLGs compared to the number of sampled individuals (MLG = 6, N = 19) (Table 1). We found low clonal richness for Panorama (R=0.08) and Asilah (R=0.28), with a few MLGs including up to 24 and 14 samples per MLG (B = 0.35 & B = 0.68). The populations in Wimereux, Roscoff, Sausset-les-pins and Calanque de Sormiou included some repeated MLGs as well. However, clonal richness was higher in the latter populations, and the Pareto B value was lower than in Panorama, Split, and Asilah (Table 1). Pairwise linkage disequilibrium (LD) was the highest in Sausset-les-pins, Calanque de Sormiou, Split, and Zahara. Multi-locus linkage disequilibrium was significant from zero in all populations but remarkably higher for Roscoff, Asilah and Panorama than other populations. This trend was the same before and after clone correction.

**Table 1.** Genetic and genotypic diversity measurements per sampled population of *Dictyota dichotoma*: The number of sampled sporophytes (N), number of multi-locus genotypes (MLG), standardised number of MLGs rarified to a sample size of 20 (eMLG), clonal richness (R), Pareto B index (B), proportion of significant linkage disequilibrium between pairs of loci (LD), multilocus linkage disequilibrium (rbarD), Observed heterozygosity (Ho), expected heterozygosity (Hs), allelic richness (Ar), inbreeding coefficient, (F_IS_), the variance of inbreeding coefficient (varF_IS_), and number of private alleles (PA). All measurements were calculated for the total dataset, and multi-locus linkage disequilibrium (rbarD), observed and expected heterozygosity (Ho and Hs), allelic richness (Ar), and inbreeding coefficient (F_IS_) were also calculated after clone-correction. Significance is indicated with * when p < 0.05.

| region | Population | N | MLG | eMLG | CR | B | LD | rbarD | Ho | Hs | Ar | $F_{IS}$ | var$F_{IS}$ | PA | ----- Clone corrected ----- | | | | |
| --- | --- | --- | --- | --- | --- | --- | --- | --- | --- | --- | --- | --- | --- | --- | --- | --- | --- | --- | --- |
|  |  |  |  |  |  |  |  |  |  |  |  |  |  |  | rbarD | Ho | Hs | Ar | Fis |
| EAtl | Goes | 61 | 61 | 20 | 1.00 | - | 0.00 | 0.04* | 0.31 | 0.35 | 2.41 | 0.15* | 0.03 | 0 | 0.04* | 0.31 | 0.35 | 1.85 | 0.15* |
| EAtl | Wimereux | 38 | 33 | 18 | 0.86 | 2.52 | 0.03 | 0.10* | 0.18 | 0.27 | 2.1 | 0.22* | 0.05 | 0 | 0.08* | 0.19 | 0.28 | 1.67 | 0.22* |
| EAtl | Roscoff | 18 | 17 | 17 | 0.94 | 4.09 | 0.24 | 0.46* | 0.64 | 0.52 | 2.96 | -0.24* | 0.09 | 0 | 0.44* | 0.64 | 0.52 | 2.35 | -0.22* |
| EAtl | Muxia | 20 | 20 | 20 | 1.00 | - | 0.03 | 0.28* | 0.44 | 0.62 | 4.61 | 0.33* | 0.07 | 4 | 0.28* | 0.44 | 0.62 | 2.87 | 0.33* |
| StroG | Zahara | 24 | 24 | 20 | 1.00 | - | 0.27 | 0.29* | 0.45 | 0.63 | 5.15 | 0.32* | 0.08 | 7 | 0.29* | 0.45 | 0.63 | 3.07 | 0.32* |
| StroG | Cap Spartel | 13 | 13 | 20 | 1.00 | - | 0.00 | 0.12* | 0.61 | 0.67 | 4.58 | 0.07 | 0.03 | 0 | 0.12* | 0.61 | 0.67 | 3.03 | 0.07 |
| StroG | Asilah | 19 | 6 | - | 0.28 | 0.68 | 0.00 | 0.67* | 0.46 | 0.34 | 2.64 | -0.17 | 0.37 | 0 | 0.34* | 0.46 | 0.51 | 2.43 | 0.08 |
| NWMed | Banyuls | 23 | 23 | 20 | 1.00 | - | 0.03 | 0.12* | 0.52 | 0.62 | 4.57 | 0.18* | 0.04 | 3 | 0.12* | 0.52 | 0.62 | 2.83 | 0.18* |
| NWMed | Girona | 20 | 20 | 20 | 1.00 | - | 0.00 | 0.08* | 0.36 | 0.56 | 3.68 | 0.34* | 0.05 | 2 | 0.08* | 0.36 | 0.56 | 2.6 | 0.34* |
| NWMed | Sausset-I.-P. | 53 | 50 | 19 | 0.94 | 3.68 | 0.59 | 0.13* | 0.63 | 0.62 | 5.03 | -0.02 | 0.02 | 1 | 0.11* | 0.63 | 0.62 | 3.04 | -0.01 |
| NWMed | Cal. d. Sormiou | 46 | 46 | 20 | 1.00 | - | 0.50 | 0.10* | 0.64 | 0.68 | 5.68 | 0.08 | 0.02 | 5 | 0.10* | 0.64 | 0.68 | 3.26 | 0.08 |
| AdSea | Split | 25 | 15 | 13 | 0.58 | 1.23 | 0.26 | 0.24* | 0.59 | 0.48 | 2.71 | -0.20* | 0.12 | 1 | 0.20* | 0.53 | 0.49 | 2.23 | -0.1 |
| EMed | Panorama | 26 | 3 | 3 | 0.08 | 0.35 | - | 1.00* | 0.63 | 0.34 | 1.92 | -0.60* | 0.46 | 0 | 1.00* | 0.58 | 0.49 | 2.12 | -0.19 |
| Mac | Punta d. Hid. | 27 | 27 | 20 | 1.00 | - | 0.14 | 0.27* | 0.42 | 0.66 | 4.61 | 0.36* | 0.03 | 8 | 0.27* | 0.42 | 0.66 | 2.96 | 0.36* |
| Total |  | 413 | 357 |  |  |  |  |  |  |  |  |  |  |  |  |  |  |  |  |

Sausset-I.-P. = Sausset-les-Pins, Cal. d. Sormiou = Callanque de Sormiou, Punta d. Hid. = Punta del Hidalgo, EAtl = European Atlantic, StroG= Strait of Gibraltar, NWMed = Northwestern Mediterranean, AdSea = Adriatic Sea, EMed = Eastern Mediterranean, Mac = Macaronesia

Before clone correction, there was a significant departure from panmixia for Panorama, Split and Roscoff, indicated by a significant, negative average F_IS_ (Table 1). Yet, distributions of the *per* locus F_IS_ indicated more negative values for Panorama, Split, Roscoff, and also Asilah (Fig. S2). The variance of F_IS_ was higher for Panorama, Asilah and Split than all other populations (Table 1). Significant homozygote excesses (positive average F_IS_) were apparent for Goes, Wimereux, Muxia, Zahara, Banyuls, Girona and Punta del Hidalgo (Table 1). After clone-censoring, a significant departure from Hardy Weinberg equilibrium (HWE) due to heterozygote excesses was only noted for Roscoff. Significant homozygote excesses after clone correction were the same as before clone correction. Allelic richness was highest in the southern populations (Calanque de Sormiou, Sausset-les-pins, Zahara, Punta del Hidalgo, Muxia, Cap Spartel) and declined both towards the northern (Wimereux and Goes) and eastern (Split and Panorama) (Table 1). The highest number of private alleles was found in Punta del Hidalgo, Zahara, Muxia, Banyuls and Calanque de Sormiou (Table 1). In the northern populations of Goes, Wimereux and Roscoff, no private alleles were observed, as in the edge populations of Panorama and Asilah.

Taken together, the low clonal richness for Panorama, Split and Asilah, combined with the negative average F_IS_ (yet non-significant for Asilah), the higher F_IS_ variance and the relatively high significant multi-locus linkage disequilibrium, indicate the high prevalence of clonality in these populations (Stoeckel et al., 2021). As for cases where a negative F_IS_ and a significant high linkage disequilibrium contradict the high clonal richness, such as observed for Roscoff, it is advised to prioritise genetic over genotypic indices (Stoeckel et al., 2021) (Table 1). Therefore, we conclude that some clonality occurs in Roscoff as well. As partial clonality will impact genotypic composition more than the genetic composition of a population, we conclude that there is also limited clonality balanced with sexual reproduction in Wimereux and Sausset-les-Pins. This would be in line with the high percentage of loci in linkage disequilibrium observed in Sausset-les-Pins. We did not observe clear patterns of clonality for the other population in Marseille (Calanque de Sormiou) based on these diversity indices, yet the percentage of loci in linkage disequilibrium is high, as one would expect in the case of clonality (Table 1).

Average yearly temperature showed a significant positive correlation with allelic richness and expected heterozygosity while a negative correlation was observed between average yearly temperature range and, allelic richness and expected heterozygosity (Table 2). Clonal richness and variance of F_IS_ did not correlate with environmental variables.

**Table 2:**
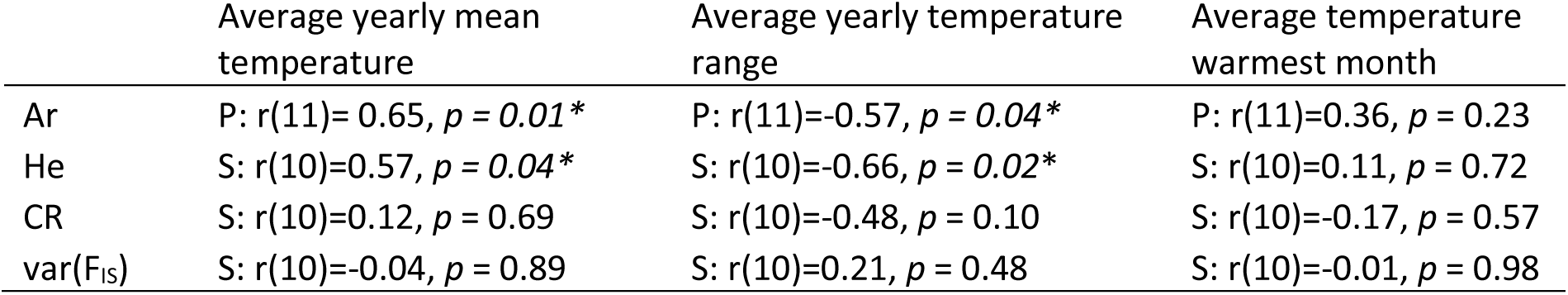
Spearman rank (S) and Pearson correlation (P) test results among genetic and genotypic diversity indices of *Dictyota dichotoma* and average yearly temperature, average yearly temperature range, and average temperature of warmest month. Ar = allelic richness, He = expected heterozygosity, CR =clonal richness, Var(F_IS_) = variance of per locus F_IS_.

### Fine-scale clonal and spatial structure

On a total of 223 sporophytes (no gametophytes found), we identified 167 unique multi-locus genotypes over the 48 quadrats scattered over the three locations in the Marseille region. Eight MLGs were repeated among quadrats within locations. In Calanque-de-Sormiou, 61 individuals were assigned to 55 MLGs scattered over 16 quadrats (Fig. 2A). One repeated MLG (MLG123) was present in 5 quadrats with a maximum of more than 100m between samples, one repeated MLG (MLG45) occurred in two quadrats in samples more than 12m apart, and another MLG (MLG44) was repeated within the same quadrat, samples 10 cm apart (Fig. 2A). In Vallon-des-Auffes, 72 individuals were assigned to 67 MLGs, with one repeated MLG (MLG32) identified in 5 samples scattered over three quadrats >100m apart, and one repeated MLG (MLG 139) was found in two samples >15m apart. In Sausset-les-Pins, clonality was remarkably higher than in the other locations in Marseille. We assigned 90 samples to 45 unique MLGs. Forty-nine samples belonged to one repeated MLG (MLG 49) scattered over 15 quadrats and >100m apart (Fig. 2A), MLG56 was identified in three samples spread over >100m, and MLG 50 was identified in three individuals scattered over three quadrats >100m apart (Fig. 2A).

**Figure 2.**
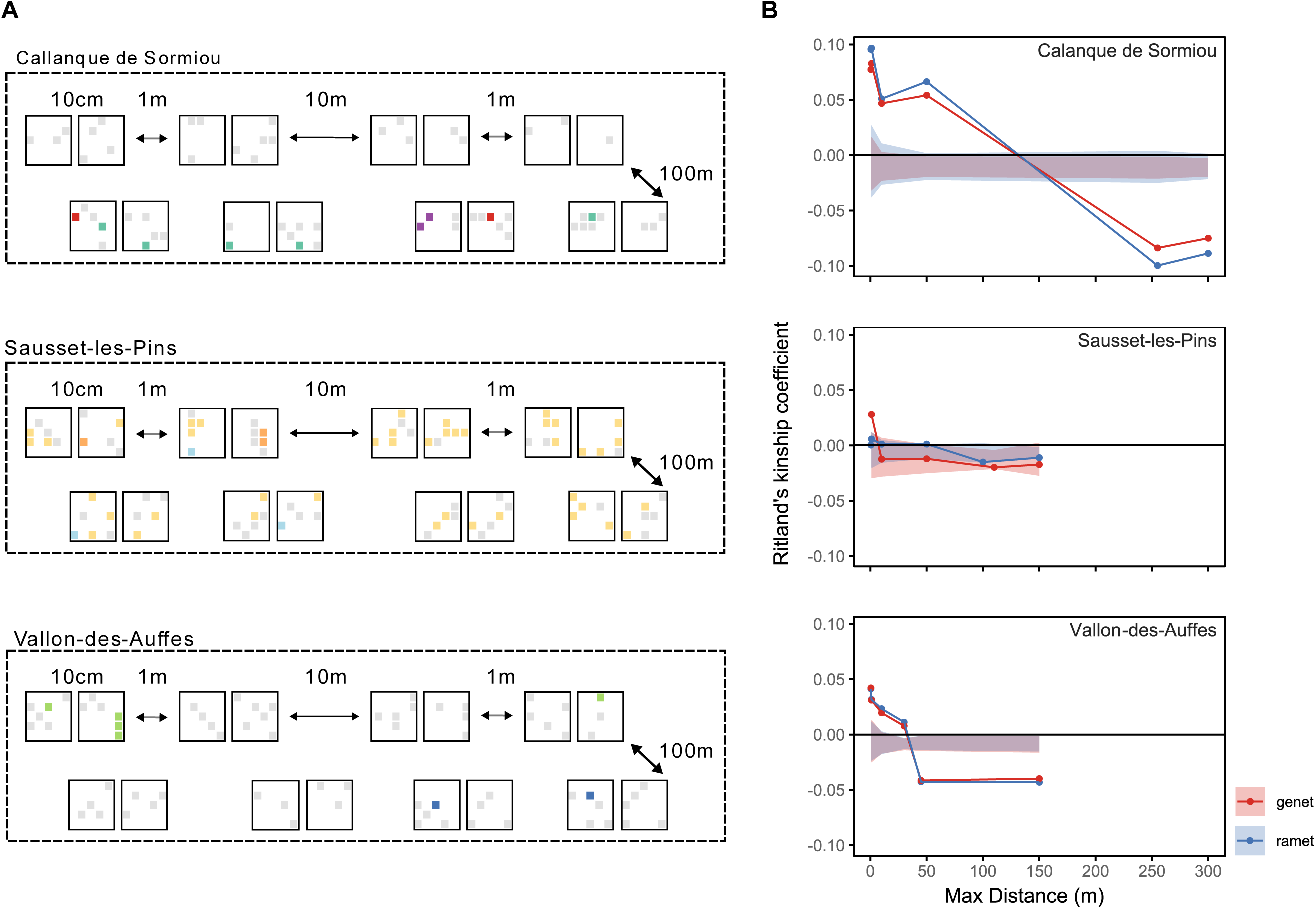
**A)** Hierarchical population structure of *Dictyota dichotoma* in Marseille region (Calanque de Sormiou, Sausset-les-Pins, Vallon-des-Auffes). Black squares represent a 50 x 50 cm sampling grid. Coloured squares indicate the MLGs of the sample, with every grey square representing a unique MLG; Other colors indicate samples belonging to a repeated MLG. **B)** Ritland’s kinship coefficient for microsatellite data for three locations within the Marseille region; Calanque de Sormiou (N = 61), Sausset-les-Pins (N = 90), Vallon-des-Auffes (N = 72) at the genet (red) and ramet (blue) level. The 95% confidence interval is indicated as a shaded area.

Ramet level spatial autocorrelation analysis indicated significant kinship coefficients for Calanque de Sormiou and Vallon-des-Auffes, with a positive signal at short distances (clustered) and a negative signal (dispersed) at larger distances. In Calanque de Sormiou, positive autocorrelation was present from 0 up to almost 150m, and when samples were more than 150m apart, autocorrelation was significantly negative (Fig. 2B). In Vallon-des-Auffes, significant positive autocorrelation reached only up to 30m and 45 and 300m the autocorrelation was significantly negative (Fig. 2B). In Sausset-les-Pins, Ritland’s kinship correlation coefficient does not deviate significantly from zero, suggesting closely related ramets are not more spatially associated as randomly expected (Fig. 2B). Genet level analysis indicated similar patterns for Calanque de Sormiou and Vallon-des-Auffes. Sausset-les-Pins indicated significant positive autocorrelation but only up to distances <10m. Starting from 50m kinship coefficients were negatively correlated with distance in Sausset-les-Pins.

### Perennial persistence vs annual recruitment

In Marseille and Girona, we collected 192 and 131 sporophytes in 2022 and 2023 and identified 161 and 131 unique MLGs, respectively (Table S3). There were 9 MLGs that were sampled in multiple grids, yet only 2 repeated MLGs were sampled over the two years, both occurring in S’Alguer, Girona (Table S3). One repeated MLG was found in the same quadrat but in 3 different grid cells (14-22 cm apart), and one repeated MLG was found in a different quadrat (approx. 8 m apart). Gametophytes were sampled in Girona only (N=28), and we found no repeated MLGs amongst the gametophytes.

When combined with the hierarchical dataset, sampled in 2021 in Marseille, we identified 4 MLGs, encompassing 54 individuals, that occurred over multiple years also in Marseille (Table 3). In Sausset-les-Pins, we sampled 42 individuals from MLG 210. In 2022, we found the same MLG in three individuals at the same site. In Calanque-de-Sormiou for MLG 199, we sampled two individuals 2021 and one in 2022 (Table 3). Four MLGs were found to occur over multiple years across sites (Table 3).

**Table 3.** The number of sporophytes and unique MLGs (in between brackets) sampled in Marseille and Girona in 2021, 2022 and 2023, and the number of individuals that belong to an MLG found over multiple years within sites (2021-2023). SLP = Sausset-les-Pins, SOR= Calanque de Sormiou, SAL= S’Alguer, FOS= Fosca.

| Location | Site | 2021 | 2022 | 2023 | 2021-2023 |
| --- | --- | --- | --- | --- | --- |
| Marseille | SLP | 90 (45) | 30 (28) | 43 (28) | 45 (1) |
| Marseille | SOR | 61 (55) | 52 (48) | 34 (25) | 3 (1) |
| Girona | SAL | - | 67 (43) | 27 (26) | 6 (2) |
| Girona | FOS | - | 45 (43) | 33 (33) | 0 |
| Total |  |  | 194 (162) | 137 (112) |  |

## Discussion

We observed differences in reproductive strategies and genetic diversity of *Dictyota* populations along its European distribution, with a clear shift in reproductive phenology marked by a sporophyte dominance and a reduction in overall fertility in Mediterranean populations driven by increased temperatures. While signals of partial clonality have been found throughout the distribution ranges, Mediterranean populations displayed a stronger signal of dominant clonality.

Environmental factors significantly influence the reproductive phenology of seaweeds, with temperature in combination with daylength being the most important drivers (de Bettignies et al., 2018; Mihaila, Magnusson, Glasson, & Lawton, 2023). Our results strongly indicate that reproductive activity aligns with seasonal temperature variations. In northern populations, which include both fertile gametophytes and sporophytes, fertility peaks when temperatures are highest. Conversely, in Mediterranean populations that are dominated by sporophytes, high temperatures appear to suppress fertility, which indicates that high temperatures (June, July) cause too much stress so that sporogenesis is inhibited. A similar trend has been observed in the temperate brown seaweed *Fucus serratus* (Fucales), where marginal populations at the southern range edge showed a significant decline in the proportion of reproductive individuals and reproductive allocation due to an increase in abiotic stress, including temperature changes (Viejo, Martínez, Arrontes, Astudillo, & Hernández, 2011). Moreover, while mean temperatures during the warmest months in the Marseille populations reached up to 24°C, *in situ* logger data indicated that temperatures exceeded 28°C on multiple days throughout summer. These real-time temperatures come close to the lethal thresholds of *Dictyota* (Delva, 2023). Indeed, macroscopic individuals are not observed during the warmest months (August, September) in Marseille, yet our results indicate that *Dictyota* can survive as filamentous or microscopic stages, allowing individuals to persist multiple years, which helps populations to endure unfavourable conditions despite reduced sexual recruitment.

Ploidy bias has been observed in multiple seaweeds and can either be sporophytes based or gametophyte based (Fierst, TerHorst, Kübler, & Dudgeon, 2005; Kain & Destombe, 1995). The dominance of one stage over the other has been hypothesised to be linked to differences in performance under certain environmental conditions. For example, diploid sporophytes of the red macroalgae *Gracillaria gracilis* (Gracilariales) performed better under high UV and high nutrient conditions than haploid gametophytes (Destombe, Godin, Nocher, Richerd, & Valero, 1993). Additionally, demographic processes such as differences in fecundity, dispersal or the survival rates of sporelings or adults might favour one ploidy level over the other (Engel, Åberg, Gaggiotti, Destombe, & Valero, 2001; Stacy A. Krueger-Hadfield & Ryan, 2020; Thornber & Gaines, 2004). However, the specific processes that drive sporophyte dominance in *Dictyota* remain to be investigated.

With respect to overall genetic diversity, a pattern of “southern richness and northern purity” is evident in our data (Hewitt, 2000). Southern populations of *Dictyota* exhibit a high proportion of endemic alleles and greater genetic diversity when considering allelic richness. This phenomenon is particularly noticeable in the populations from Punta del Hidalgo in the Canary Islands, the Iberian Peninsula (specifically Muxia and Zahara), and to a lesser extent in the western Mediterranean (including Banyuls and Calanque de Sormiou). These regions have previously been identified as potential glacial refugia for warm temperate seaweeds (Hoarau, Coyer, Veldsink, Stam, & Olsen, 2007; Maggs et al., 2008). However, heterozygosity does not show a clear trend. Allelic richness is distinct from measures of heterozygosity because it is tied to a species’ long-term evolutionary potential. Additionally, it is more sensitive to population bottlenecks and is better at reflecting changes in rare alleles (Allendorf, 1986; Greenbaum, Templeton, Zarmi, & Bar-David, 2014). The observed gradual decline in allelic richness and the absence of private alleles in northern populations support the scenario of northward colonization of areas that were previously glaciated following the last glacial maximum, indicating that also past climate conditions influence contemporary genetic diversity (Hampe & Petit, 2005; Hewitt, 2000).

Identifying asexual reproduction can be challenging since rare recombination events may erase the genetic indicators of clonality (Halkett, Simon, & Balloux, 2005; Stacy A Krueger-Hadfield, Guillemin, Destombe, Valero, & Stoeckel, 2021; Stoeckel et al., 2021). However, we identified several signs of clonality within multiple populations. These included elevated levels of heterozygosity resulting in negative Fis values, a larger variance of Fis, increased linkage disequilibrium, repeated MLGs. We found signs of limited clonality mixed with sexual reproduction in some northern populations (Wimereux, Roscoff) and more prominent clonality in southern populations (Asilah, Split, Panorama, Sausset-les-Pins). Yet, no significant link between clonality and temperature could be found. However, in this study we used satellite-derived temperature data, which may not accurately capture local-scale temperature variations (Verdura et al., 2021), or effectively represent the relationship with clonality.

Interestingly, the clonality patterns measured in Sausset-les-Pins varied significantly between the samples taken in 2015 to assess diversity across the species range and those taken in 2021 for detailed examination of clonality and spatial structure, particularly regarding genotypic indices. Sampling effort in 2015 (53 samples) indicated high genotypic richness and only limited clonality, while sampling in 2021 (90 samples) indicated much lower genotypic richness. This further supports findings of Stoeckel et al. (2021), indicating that genotypic parameters tend to be overestimated when using smaller sample sizes, which in turn leads to a substantial underestimation of clonality. Additionally, clonality patterns appear to differ a lot among sampling sites within the Marseille region. In Sausset-les-Pins, individuals belonging to the same MLG spanned more than 100m, while in Calanques-de-Sormiou and Vallon des Auffes, the maximum observed distance among clones was approximately 16 m, yet also here, sample sizes vary between 61 and 90 individuals. Altogether, these observations suggest that careful quantification of (partial) clonality is essential.

Brown seaweeds can reproduce clonally through a variety of mechanisms (Vranken et al., 2023). Our results suggest that apomeiosis is not frequent in cultured strains *Dictyota’s*. This might, however, be different for individuals in the field if clonality were to be induced by environmental variables, as other results suggest. Other mechanisms that *Dictyota* could use to reproduce clonally are fragmentation (Herren et al., 2006), sprawling or by the formation of *in-situ* germlings formed by mitotic monospores ((Hwang et al., 2005), pers obs. C. Rodríguez-Prieto). Yet more research, would be needed to identify the relative importance of the various modes of clonal reproduction. To assess clonality, we identified unique MLGs based on a psex threshold of 0.05. So, individuals resulting from fertilisation between egg and sperm coming from dioicious gametophytes produced by the same sporophyte, i.e. intergametophytic selfing (Klewoski & Edward, 1969), arise from the same sexual event. However, since they exhibit a psex value of less than 0.05, they belong to the same MLG and are considered clones in our study. Importantly, frequent intergametophytic selfing should result in a heterozygote deficiency and significant positive F_IS_ values. Since this is not observed in Sausset-les-Pins (not in 2012 and not in 2021 Table S4), and we only sporadically observed fertile gametophytes, we conclude that only very rarely intergametophytic selfing may contribute to this pattern of clonality.

We also note that given the reproductive phenology and likely the reproduction type (asexual *vs* sexual) is linked to temperature, these aspects might be further impacted by global change-related changes in seawater. Previous research indicated that temperatures in the Mediterranean would rise to levels unsuitable for *Dictyota* (Delva, 2023) if it does not adapt to higher temperatures. The southern Mediterranean populations bear higher standing genetic variation (higher allelic richness) compared to northern populations, which should benefit their evolutionary potential. However, for these populations to adapt to warmer temperatures, natural selection needs to occur, which is only efficient if sexual reproduction is possible (Hartfield, 2016). If Mediterranean *Dictyota’s* shift their phenology and move towards loss of fertility and clonal reproduction as a mechanism to survive warm temperatures, this could lead to an evolutionary dead end in the long term (Hartfield, 2016).

## Conclusion

This study demonstrates how life cycle variation in the brown macroalga *Dictyota dichotoma* is influenced by large-scale environmental gradients, particularly temperature, across its European distribution range. Our comprehensive analysis of phenology, genetic diversity, and genotypic variation reveals a significant shift from predominantly sexual reproduction in the Atlantic populations to a dominance of sporophytes and more prominent clonal reproduction in the Mediterranean populations. These shifts are linked to reduced fertility and the capacity to survive over multiple years Mediterranean populations, allowing them to persist under suboptimal thermal conditions. The observed latitudinal gradient in allelic richness and the presence of unique alleles in southern populations represent a post-glacial recolonization scenario as seen in other marine taxa. Importantly, the loss of sexual reproduction in Mediterranean populations may limit their adaptive potential in the face of climate change, raising concerns about long-term persistence in warming seas. Our findings contribute to understanding how life cycle plasticity, clonality, and regional environmental pressures interact to shape genetic structure and reproductive strategies in marine macroalgae. Integrating life cycle variation into biogeographical frameworks will be crucial for forecasting species responses to changing environments, especially in habitat-forming taxa like macroalgae that underpin coastal ecosystem function.

