## Supplemental Files for "Rare sex but a long life sustain seaweed populations at the warm edge of their range"

### Supplementary Material

**Table S1** Akaike Information Criterion (AIC) and Bayesian Information Criterion (BIC) for multiple model variations of Rodríguez et al. (2019), Brière et al. (1999), Blanchard et al. (1996) and a linear model.

| **Model** | **AIC** | **BIC** |
|---|---|---|
| Rodriguez et al. (2019) (Region specific b and e) | -40.8 | -26.1 |
| Rodriguez et al. (2019)(Fully parametrized) | -35.1 | -15 |
| Brière et al. (1999) | -29.1 | -12.6 |
| Blanchard et al. (1996) | -14.8 | 1.65 |
| Linear model | 378 | 386 |

**Table S2** Sporophyte information for which offspring of *Dictyota dichotoma* tetrads have been monitored in the laboratory. Strain info (Parent), strain ID in the Ghent University Culture Collection (GUCC), the location where the parent sporophyte has been sampled (Location), Male:female ratio, and the ploidy level of the offspring's genotype.

| **Parent** | **GUCC** | **Location** | **Offspring phenotype % male / % female** | **Offspring genotype** |
|---|---|---|---|---|
| A1C | UGCC0031 | Goes - Atl. Oc. | 55 / 45 (n=40) | 100% haploid (n= 32) |
| Mar DII | UGCC0023 | Marseille - Med. Sea | 43 / 57 (n=7) | 100% haploid (n=8) |
| Mar15C | UGCC0019 | Marseille - Med. Sea | 56 / 44 (n=16) | 100% haploid (n=24) |
| Dd Croatia | UGCC0003 | Croatia – Med Sea | 50 / 50 (n=24) | 100% haploid (n=23) |

**Table S3** The number of sporophytes, gametophytes and unique MLGs (in brackets) sampled in Marseille and Girona in 2021, 2022 and 2023, and the number of individuals that belong to an MLG found over multiple years within sites (2021-2023). N= number sampled individuals, MLG = number of unique multilocus genotypes. MAR = Marseille, GIR = Girona, SLP = Sausset-les-Pins, SOR= Calanque de Sormiou, VAL= Vallon-des-Auffes, SAL= S'Alguer, FOS= Fosca.

|  |  |  |  | ---------------------- sporophytes ---------------------- |  |  | ---------------------- gametophytes ---------------------- |  |  |
|---|---|---|---|---|---|---|---|---|---|
| Location | Site | Month | Quadrat | N (MLG) 2022 | N (MLG) 2023 | MLG 2022+2023 | N (MLG) 2022 | N (MLG) 2023 | MLG 2022+2023 |
| MAR | SLP | June | Q1 | 3 (3) | 8 (8) | 0 | 0 | 0 | - |
| MAR | SLP | June | Q2 | 6 (6) | 4 (4) | 0 | 0 | 0 | - |
| MAR | SLP | June | Q3 | 21 (20) | 13 (11) | 0 | 0 | 0 | - |
| MAR | SLP | June | Q4 | - | 18 (7) | 0 | 0 | 0 | - |
| MAR | SOR | June | Q1 | 9 (9) | 5 (5) | 0 | 0 | 0 | - |
| MAR | SOR | June | Q2 | 21 (20) | 0 | 0 | 0 | 0 | - |
| MAR | SOR | June | Q3 | 14 (11) | 4 (4) | 0 | 0 | 0 | - |
| MAR | SOR | Nov | Q3 | 6 (6) | 0 | 0 | 0 | 0 | - |
| MAR | SOR | June | Q2a | - | 10 (9) | 0 | 0 | 0 | - |
| MAR | SOR | June | Q2b | - | 9 (8) | 0 | 0 | 0 | - |
| GIR | SAL | June | Q1 | 31 (25) | 7 (7) | 0 | 1 (1) | 1 (1) | 0 |
| GIR | SAL | June | Q2 | 36 (19) | 20 (19) | 6 (2) | 1 (1) | 0 | - |
| GIR | FOS | June | Q1 | 16 (14) | 19 (19) | 0 | 6 (6) | 3 (3) | 0 |
| GIR | FOS | June | Q2 | 12 (12) | 5 (5) | 0 | 7 (7) | 1 (1) | 0 |
| GIR | FOS | June | Q3 | 17 (17) | 9 (9) | 0 | 7 (7) | 1 (1) | 0 |
| Total | | | | 192 (161) | 131 (109) | | 22 (22) | 6 (6) | 0 |

**Table S4** Genetic and genotypic diversity measurements per for *Dictyota dichotoma* sampled in the region of Marseille, France: The number of sampled sporophytes (N), number of multi-locus genotypes (MLG), clonal richness (R), Pareto B index (B), multilocus linkage disequilibrium (rbarD), Observed heterozygosity (Ho), expected heterozygosity (Hs), allelic richness (Ar), inbreeding coefficient, ($F_{IS}$), the variance of inbreeding coefficient (var$F_{IS}$), and number of private alleles (PA). Significance is indicated with * when $p < 0.05$. .

| Population | N | MLG | R | B | rbarD | Ho | Hs | Ar | $F_{IS}$ | var$F_{IS}$ | PA |
|---|---|---|---|---|---|---|---|---|---|---|---|
| Calanque de Sormiou. | 61 | 55 | 0.90 | 2.42 | 0.24* | 0.42 | 0.55 | 6.57 | 0.19* | 0.04 | 12 |
| Sausset-les-Pins | 90 | 45 | 0.49 | 0.91 | 0.57* | 0.58 | 0.50 | 5.78 | -0.09 | 0.10 | 3 |
| Vallon-des-Auffes | 72 | 67 | 0.93 | 2.52 | 0.11* | 0.51 | 0.60 | 6.53 | 0.11* | 0.04 | 4 |

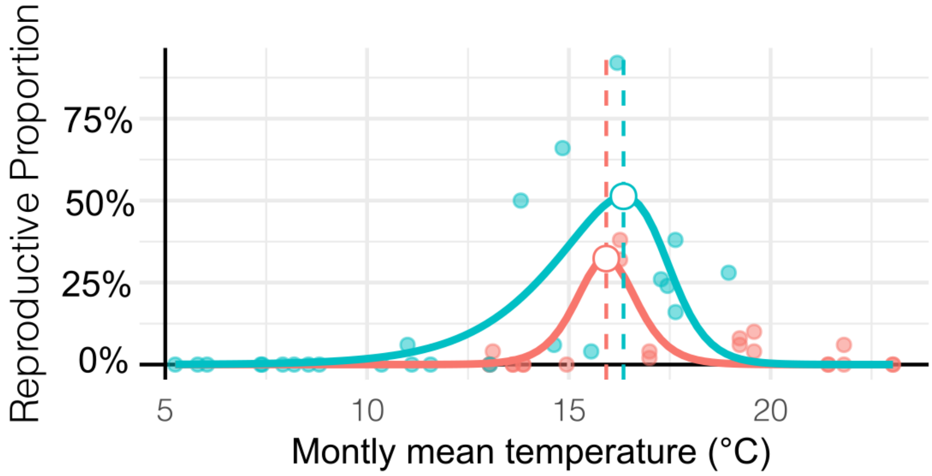


**Figure S1:** The proportion of reproductive sporophytes and gametophytes (fertility) of Dictyota dichotoma per monthly mean temperature (°C) modelled using the model of Rodríguez et al. (2019). In the selected Rodriguez model ((with Region specific b and e) (p, peak fertility is predicted to occur at 15.9 °C for Mediterranean populations and 16.4 °C for North Sea populations. Mediterranean Sea (red) includes samples from Sausset-les-pins and Callanque de Sormiou and North sea populations (blue) includes samples from Goes and  Wimereux.

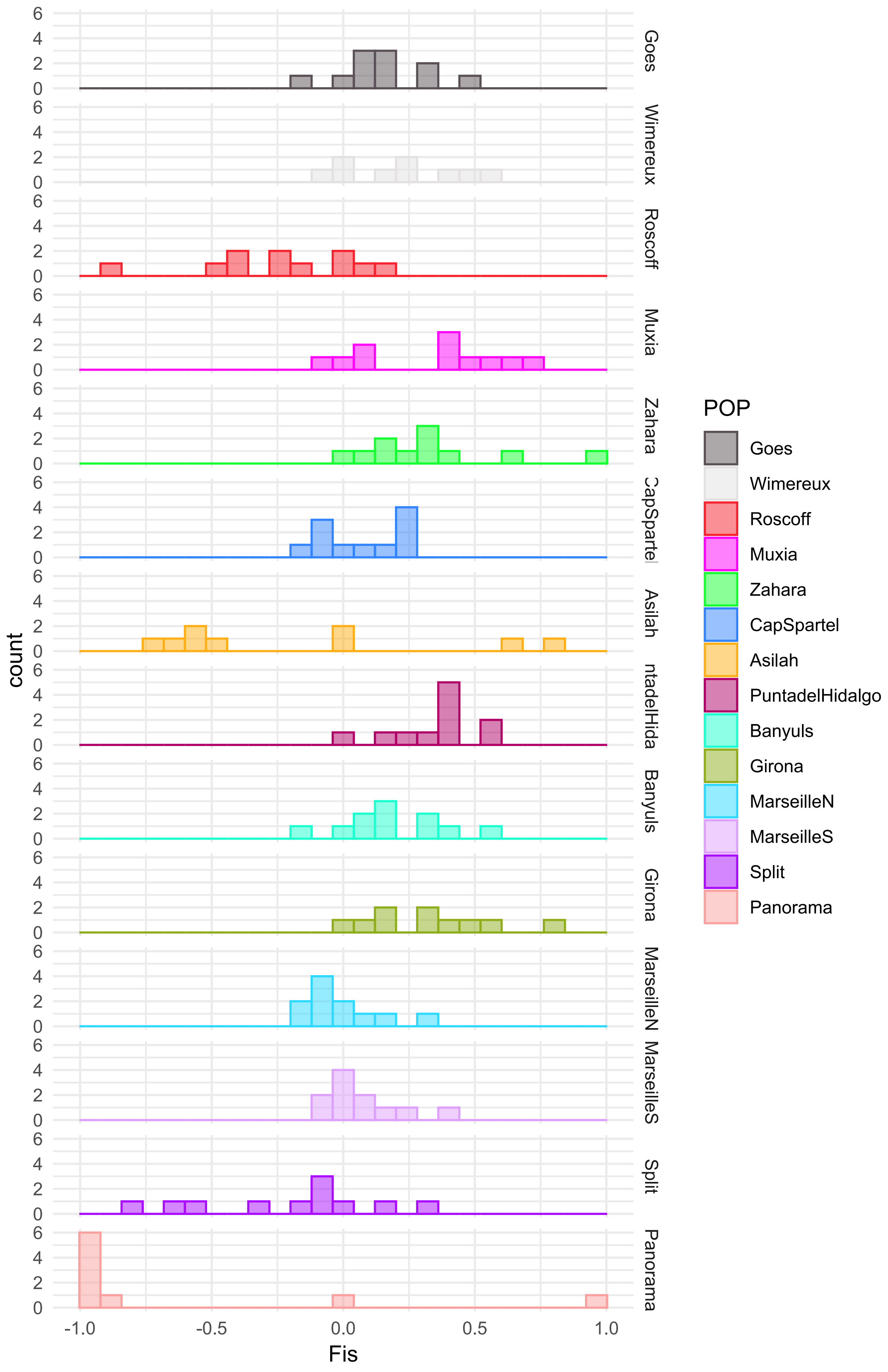


**Figure S2**: Per locus $F_{IS}$ distribution for every population calculated before clone correction
